# Structural basis for heat tolerance in plant NLR immune receptors

**DOI:** 10.64898/2025.12.17.694812

**Authors:** Marta Grech-Baran, Kamil Witek, Hee-Kyung Ahn, Małgorzata Lichocka, Teresa Vargas-Cortez, Izabela Barymow-Filoniuk, Agnieszka I. Witek, Jacek Hennig, Jonathan D. G. Jones, Jarosław T. Poznański

**Author notes:** To whom correspondence should be directed E-mail: Marta Grech-Baran.

## Abstract

Nucleotide-binding leucine-rich repeat (NLR) immune receptors sense pathogen molecules and oligomerize, initiating defense signaling. Some NLRs function poorly at elevated temperatures for unknown reasons. We show that temperature-sensitive NLRs retain ligand binding at elevated temperatures but are impaired in oligomerization. We identify key residues involved in temperature resilience. Structural modeling reveals stabilizing intramolecular interactions of the NB-ARC domain with surface residues of the adjacent leucine-rich repeat (LRR) that preserve receptor integrity and functionality under heat stress. These insights enable *in silico* classification of NLRs as temperature-sensitive or -tolerant and underpin design of temperature tolerant variants of temperature sensitive NLRs.

These findings provide a mechanistic basis for temperature sensitivity in plant immune receptors and enable engineering of temperature-tolerant disease resistance in crops.

## Introduction

The climate emergency threatens global agriculture and food security. Elevated temperatures weaken plant immune responses by inactivating immune receptors or blocking systemic acquired resistance (*1*), yet the molecular mechanisms remain largely unknown (*2*). Plants employ two main classes of immune receptors: cell-surface pattern recognition receptors (PRRs) and intracellular nucleotide-binding leucine-rich repeat receptors (NLRs) (*3*). PRRs detect pathogen-associated molecular patterns to trigger pattern-triggered immunity (PTI), whereas NLRs directly or indirectly perceive pathogen effectors (AVRs) to initiate effector-triggered immunity (ETI) (*4*). Within the NLR network, sensor NLRs recognize effectors, while helper NLRs act downstream to execute defense (*5*).

An NLR consists of an N-terminal signaling domain (either Coiled-Coil (CC), CC-RPW8 (CCR), or Toll/Interleukin-1 receptor (TIR)), a conserved nucleotide-binding and oligomerization NB-ARC domain, and a C-terminal Leucine-Rich Repeat (LRR) domain. The NB-ARC module comprises three subdomains: the nucleotide-binding domain (NBD), helical domain 1 (HD1), and winged-helix domain (WHD). This module functions as a molecular switch controlling NLR oligomerization (*6, 3, 7, 8*). In both CC-NLRs (CNLs) such as ZAR1 and TIR-NLRs (TNLs) such as Roq1 and RPP1, the WHD region is critical for oligomerization, stabilizing inter-subunit contacts within the resistosome complex (*9, 10, 11*). Upon effector recognition, several CNLs and CCR-NLRs assemble into cation channels that drive Ca²⁺ influx and hypersensitive response (HR) (*12*). ZAR1 and Sr35 form pentameric resistosomes (9, 13), NRC2 and NRC4 assemble as hexamers (*14, 15*), and TNLs such as Roq1 and RPP1 form tetramers with NADase activity to generate signaling messengers (*10, 11, 16*).

Temperature sensitivity of plant immunity was first noted when resistance to tobacco mosaic virus (TMV) mediated by the *N* gene was lost as temperature increased from 22°C to 28°C (*17*). Similarly, the tomato nematode resistance genes *Mi-1.1* and *Mi-1.2* are inactivated by heat (*18*), and PVY resistance conferred by *Ny-1* is abolished at elevated temperature (*19*). In contrast, some stem rust resistance genes such as *Sr21* and *Sr13* remain functional or even stronger at elevated temperatures, while *Sr6*, *Sr10*, *Sr15*, and *Sr17* lose activity (*20, 21*). These observations suggest that temperature affects NLR activation, which requires extensive conformational rearrangements and structural stabilization. Elevated temperature is known to disrupt secondary and tertiary protein structures, potentially impairing oligomerization (*22*).

*Ry_sto_* is a TNL conferring resistance to PVY and related potyviruses (*23*). Unlike many NLRs, *Ry_sto_* retains function above 30°C (*23, 24*). Here we identify the molecular mechanism underlying its temperature tolerance through structural modeling and functional assays. We show that this mechanism is generalizable across NLRs, including helper types, and propose an *in silico* framework for predicting and engineering temperature-tolerant immune receptors.

## Results

### 1.1 Oligomerization or ligand binding: what fails at elevated temperatures?

To better understand temperature tolerance of Ry_sto_, we determined whether elevated temperature impacts the receptor’s oligomerization or its binding to AVR. As a comparator, we used the N receptor against TMV, which exhibits temperature sensitivity at 28°C (*17*). Following transient co-expression of Ry_sto_ and N with their respective ligands (PVY Coat Protein (PVY-CP) and TMV p50) at permissive and elevated temperature, we performed Blue Native PAGE (BNP) that allows detection of native protein complexes (*25*). At 22°C, both N and Ry_sto_ formed high-molecular-weight oligomeric complexes that migrated above 720 kDa (Fig. 1A). At 28°C, Ry_sto_ still oligomerized strongly, while only traces of oligomers were detected for N, both for extract and Co-IP-ed samples (Fig. 1A, B). Surprisingly, both Ry_sto_ and N receptors interacted with their ligands at each temperature tested (Fig. 1C, D). This suggests that temperature insensitivity of oligomerization is crucial for temperature tolerance.

**Fig. 1.**
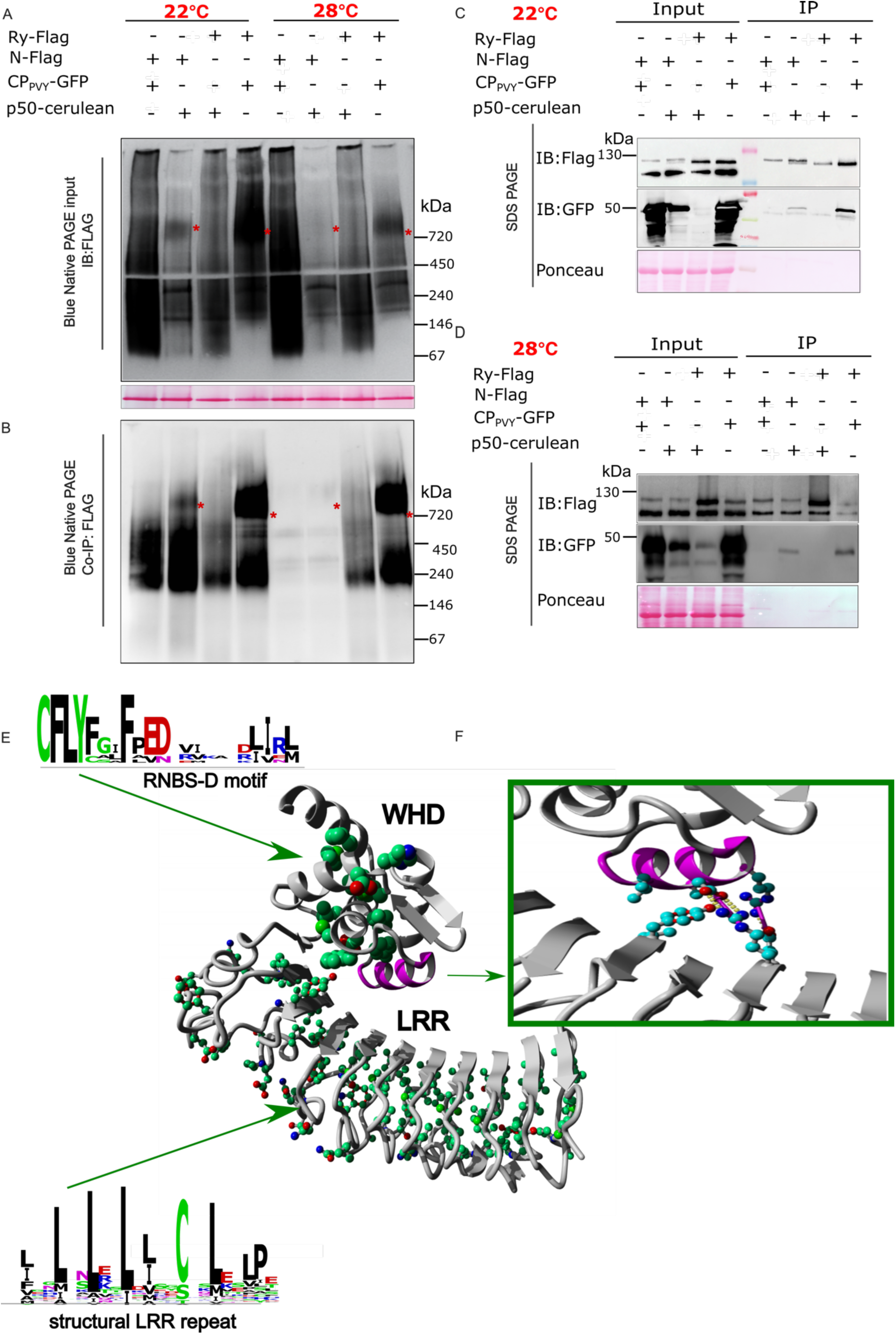
Contribution of oligomerization and stabilization of the tertiary structure of NB domain to temperature tolerance of R receptors. Temperature-sensitive (TS) and temperature-tolerant (TT) receptors show different patterns of oligomerization after AVR-dependent activation at permissive (22°C) and elevated (28°C) temperature. Protein lysates from the leaves of *eds1* knockout *N. benthamiana* co-infiltrated with Flag-tagged Ry_sto_ (TT) or N (TS) and their respective ligands (PVY-CP and p50) at each temperature, were purified using anti-Flag beads. Input samples (A) and immunoprecipitated proteins eluted with 3xFlag peptide (B) were loaded onto Blue Native PAGE (BNP). N-Flag co-expressed with PVY-CP-GFP and Ry_sto_ with p50-cerulean were used as controls. SDS-boiled input and IP eluates from 22°C (C) and 28°C (D) were loaded onto SDS–PAGE and labelled with anti-Flag or anti-GFP antibodies. Ponceau S staining served as a loading control. Data information: At least three biological replicates and corresponding technical replicates were tested. (E) A representative structure of the WHD region of NB domain together with the LRR domain of a R receptor. Most WHD-LRR contacts initiate downstream from the divergent RNBS-D motif CFbYC/FxxFPED (green). The divergent residues form the third helix (magenta), which is crucial for maintaining the conformation of the region through interactions with the LRR domain. Green balls – carbon, red – oxygen, blue – nitrogen atoms. Stretches of axxa/Lxxa/LxxL/Ixa/LxxCxxa/Lxxaxx motifs forming the LRR strands are marked in green. (F) Zoom-in on the third helix with WHD-LRR interfacing occurring between beta strands 3 to 6 of the LRR. Favorable electrostatic interactions (salt bridges and hydrogen bonds) are indicated with arrows.

### 1.2 Mapping the temperature-tolerant region of NLR receptors

The WHD subdomain of the NOD domain undergoes marked conformational rearrangements during resistosome assembly, which may be influenced by elevated temperature (*8, 10, 22*). The five α-helices of the WHD interact with the HD1 region of the NOD and LRR domains. Within MHD module, the conserved MHD motif in the fourth helix is essential for ADP binding, whereas the divergent RNBS-D motif (CFbYC/FxxFPED) in the third helix contributes to structural stability through intramolecular contacts with the LRR domain (Fig. 1E, F). During oligomerization, the third helix positions the fourth helix such that the MHD region rotates away from the nucleotide-binding site, exposing the oligomerization interface (*12*). In Ry_sto_, modeling identified three types of intramolecular interactions, hydrophobic contacts, hydrogen bonds, and salt bridges, that stabilize the WHD relative to the multistranded LRR motif. These include E456 (WHD)–K624 (LRR), D450 (WHD)–H625 (LRR), and I453 (WHD)–K624 (LRR) (Fig. 2A). Such interactions likely reinforce the tetrameric structure and contribute to adaptation under variable environmental conditions.

**Fig. 2.**
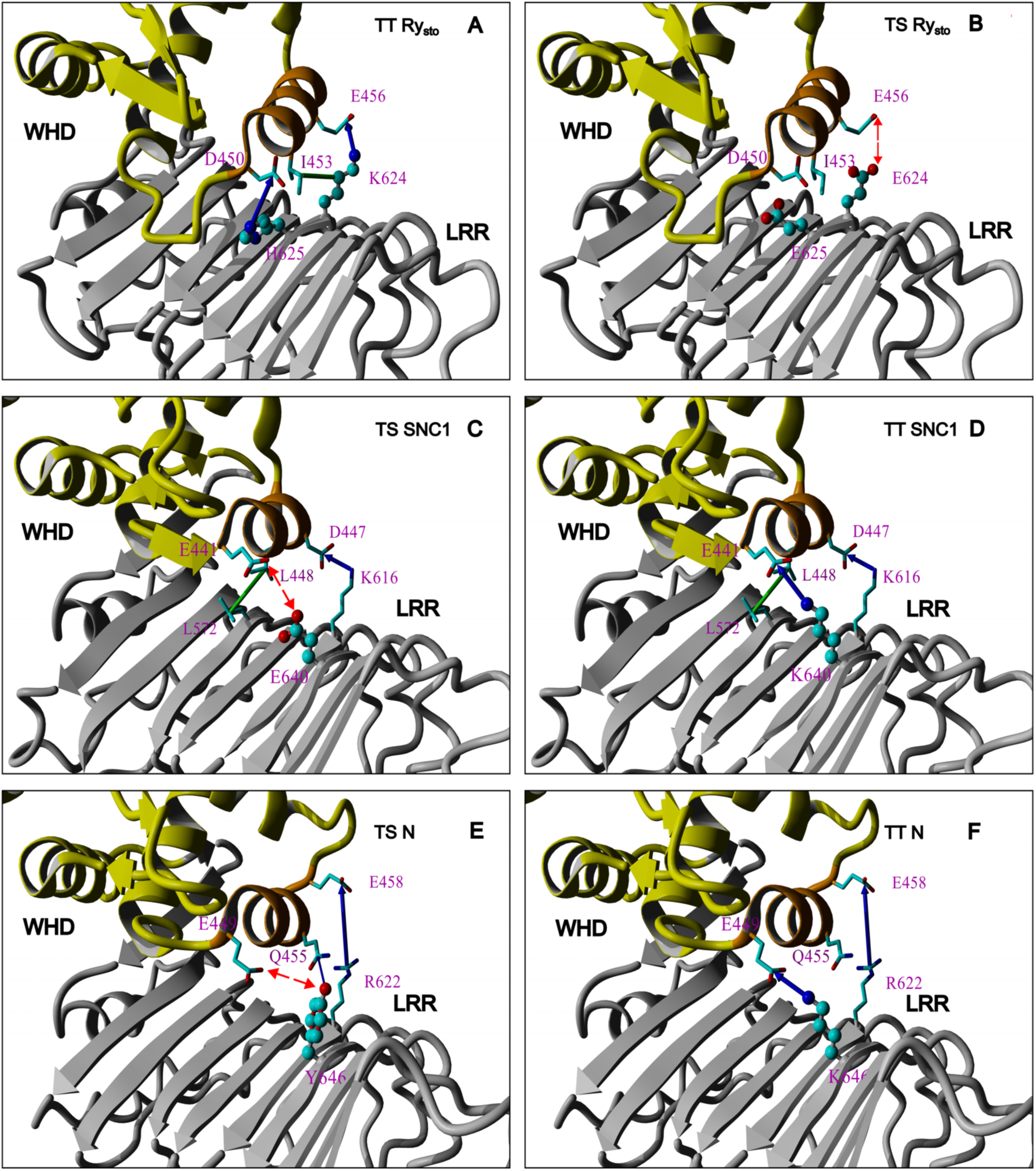
Predicted WHD-LRR interactions in TS and TT R receptors and their variants. (A) TT Ry_sto_ and (B) the predicted TS form of Ry_sto_ with reduced number of contacts; (C) native TS SNC1 and (D) its TT variant, with new stabilizing surface contacts; (E) native N receptor and (F) TT N variant, with increased number of favorable interactions. Yellow – WHD, gray – LRR, gold – the interacting third helix. The crucial residues are named and shown as stick or ball-and-stick (when mutated) cartoons. Green balls – carbon, red – oxygen, blue – nitrogen atoms. Blue arrows indicate favorable electrostatic interactions (salt bridges and hydrogen bonds), red arrows show the unfavorable ones, and green arrows represent hydrophobic contacts. Confidence predictions for the modeled regions are provided in Suppl. Table 2.

Two TNL receptors, SNC1 and N, are intrinsically temperature-sensitive (TS) but there exist temperature-tolerant (TT) variants of each (*26*). Sequence polymorphisms associated with TT variants of each were discovered, but no structural generality was derived from these findings (*26*). In TT SNC1, our modeling suggests that the E640K (LRR) substitution restores resistance at 28°C by forming a new stabilizing salt bridge with E441 (WHD). In contrast, the wild-type (WT) TS SNC1 maintains a D447 (WHD)–K616 (LRR) contact and weaker hydrophobic interactions such as L448 (WHD)–L572 (LRR) (Fig. 2C, D). Similarly, the TS variant of N is modelled to form E458 (WHD)–R622 (LRR) and Q455 (WHD)–Y646 (LRR) interactions, with the Y646K (LRR) mutation likely introducing an additional ionic bond that enhances stability (Fig. 2E, F).

Comparative modeling across more than forty receptors (Suppl. Fig. 1; Suppl. Table 1) revealed that TT variants exhibit denser WHD–LRR contact networks. In TT Rx, E426 (WHD)–R628 (LRR) and K406 (WHD)–D543 (LRR) interactions promote compactness through supplementary electrostatic contacts involving the fourth helix. TT Sr21 is stabilized by hydrogen bonds and hydrophobic contacts (Suppl. Fig. 2B), whereas the TS receptor Mi1.1 shows markedly fewer such interactions (Suppl. Fig. 2C). Distinct patterns of WHD–LRR interaction were observed between TT and TS receptors. In the WHD-LRR interaction, contacts primarily involve the third WHD α-helix and the termini of the second and fourth helices, particularly near the divergent region downstream of the RNBS-D motif (Fig. 1E). The corresponding LRR interface spans β-strands 3–6, comprising 23-amino-acid repeats that form stable motifs (axxaxxa/LxxL/LxaxxCxaxxax). These repeats influence WHD conformation through intra- and intermolecular interactions, while conserved residues within the LRR core remain buried and do not directly participate in WHD–LRR contacts.

We further examined receptors from high-temperature environments, including Yr87/Lr85 (Mediterranean) and Yr5a/Yr7 (China, India, and Australia), which confer resistance to various rust pathogens. Yr5a and Yr7 display strong electrostatic interactions E608 (WHD)–R797 (LRR) and E599 (WHD)–R818 (LRR), as well as intramolecular contacts with the terminus of the fourth WHD helix (Suppl. Fig. 3A, B). In Yr87/Lr85, stability is achieved through multiple interdomain hydrogen bonds (Q450 (WHD)–D584 (LRR), Y432 (WHD)–Q450 (WHD), R446 (WHD)–D618 (LRR), N452 (WHD)–E638 (LRR)) and extensive hydrophobic interactions (Suppl. Fig. 3C). Collectively, these results suggest that temperature sensitivity in NLR receptors is governed by the structural integrity of the WHD–LRR interface, with enhanced interdomain stabilization underlying temperature tolerance.

### 1.3 *In silico* classification of NLR receptors by their predicted temperature sensitivity

To determine the TT or TS status of NLR receptors *in silico*, we applied three rules. First, at least three electrostatic interactions are required between the third alpha helix of the WHD, and the beta strands on the LRR surface. Second, stabilization between the WHD region and the LRR domain is enhanced when hydrogen bonds or charge-charge interactions stabilize the orientation of both termini of the third alpha helix of WHD relative to the spatially interacting region of the beta sheet of LRR. Third, we consider the additional stabilization occurring in the surrounding regions (termini of the second and fourth helix) by intramolecular interactions with the HD1 region.

We applied these rules to a set of immune receptors of which temperature sensitivity status was either previously established (**bold**) or unknown. The classification yielded: TT – **Rx**, **Ry_sto_**, Rpi-abpt1, Rpi-amr3, Gpa2, Rpi-chc1, Rpi-edn1.1, NRG1, NRC2, NRC3, NRC4, **Sr21**, BS4; semi-TT – Rpi-amr1, Hero, Rp1, Ph3, Roq1, PvR4, RP1, R1, Rx2, **Tm2**, Rpi-blb3, Rpi-mcq1; and TS – Gro1-4, I2, **Mi-1.1**, **Mi-1.2,** R2, R3a, R3b, R8, Rpi-bt1, Bs2, Rpi-blb1, Rpi-blb2, Rpi-sto1, Y-1, L4, **N, SNC1, N’**. All previously experimentally classified receptors fell into the correct categories, further increasing confidence in the proposed model and supporting the classification pipeline. The proposed models, criteria, and specific interactions for each receptor are listed in Suppl. Fig. 1 and Suppl. Table 1, respectively.

### 1.4 Validation of the model by reversing temperature status or Ry_sto_ and Roq1

As predicted, at least three WHD–LRR interactions were essential for maintaining Ry_sto_ function at elevated temperatures (Fig. 2A). To test this, we used Yasara template-based homology modeling combined with AI (*27*) to design GFP-tagged putative temperature-sensitive mutants, Ry_sto-K624E_ and Ry_sto-H625E_ (Fig. 2B). These and the WT Ry_sto_-GFP were transiently co-expressed with Strep-tagged PVY-CP (AVR) at permissive (22°C) or elevated (30°C) temperature. At 22°C, all variants triggered comparable effector-dependent hypersensitive responses (HR) and ion leakage (Fig. 3A–C). At 30°C, however, HR was completely lost in Ry_sto-K624E_ but remained strong in the WT Ry_sto_ and Ry_sto-H625E_ (Fig. 3D–F), indicating that residue K624 is critical for temperature tolerance.

**Fig. 3.**
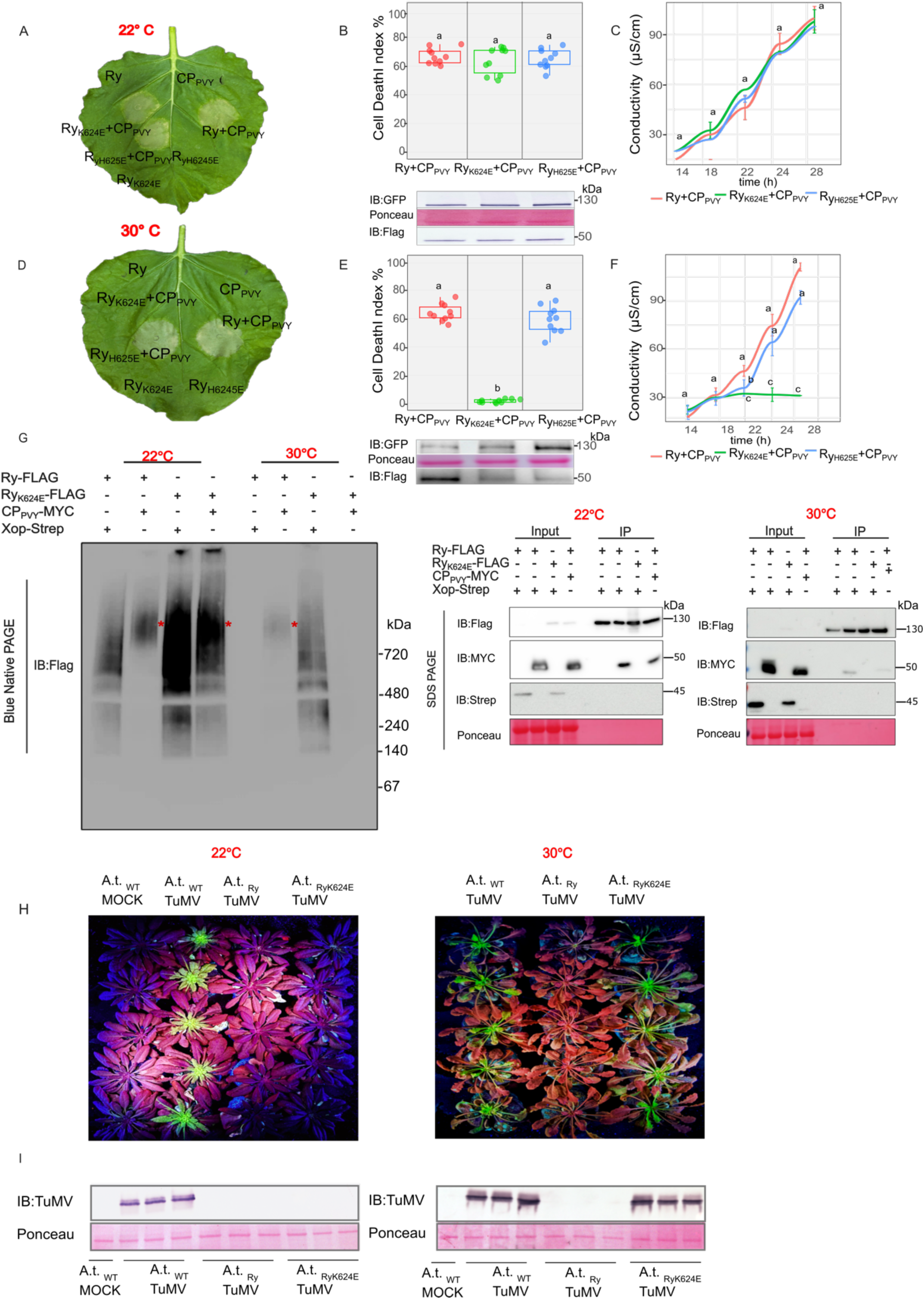
Ry_sto_ temperature tolerance is directly influenced by the stabilization of the WHD-LRR region that facilitates oligomerization at elevated temperatures. Ry_sto_ variants were co-infiltrated with PVY-CP into *N. benthamiana* leaves at permissive (22°C; A-C) or elevated (30°C; D-F) temperatures. Representative HR phenotypes (A, D), HR scoring box plots and protein accumulation of Ry_sto_ variants (B, E), and ion leakage measurements (C, F) are shown for each temperature. Target bands were immunoblotted with appropriate antibodies, with Ponceau S staining serving as a loading control. Protein extraction and antibody detection were performed at 3 dpi, while HR was quantified and documented at 5 dpi. Data information: At least three biological replicates and corresponding technical replicates were tested. (G) TS variant of Ry_sto_ oligomerizes at permissive, but not at elevated temperature. Proteins were extracted from the leaves of *N. benthamiana eds1* knockout plants co-expressing Ry_sto_-Flag or the mutated Ry_sto-K624E_-Flag with PVY-CP-Myc, or with Xop-Strep as a control, at permissive and elevated temperatures. Extracts from both temperatures were aliquoted and loaded onto BNP gels, followed by immunoblotting using an anti-Flag antibody, or co-immunoprecipitated with anti-Flag beads and eluted using Flag peptide. The eluates were subsequently incubated in 3xSDS sample buffer with DTT, loaded onto SDS-PAGE and subjected to immunoblotting with anti-Flag, anti-Myc, or anti-Strep antibodies, respectively. Data information: At least three biological replicates and the corresponding technical replicates were tested. (H) TS form of Ry_sto_ confers systemic resistance to TuMV infection at permissive but not at elevated temperature. Transgenic wild-type (WT) Ry_sto_ and Ry_sto-K624E_ Arabidopsis lines, and WT control plants, were infected with TuMV-GFP through agroinfiltration at both temperatures. The viral spread was monitored by GFP fluorescence at 14 dpi. Photographs were taken under blue light (488 nm). SDS-PAGE for protein samples from permissive and elevated temperature (I) was performed at 14 dpi. Immunoblotting was performed using anti-TuMV antibodies. Ponceau S staining served as a loading control. Data information: At least 75 plants from five different lines of each variant were tested, with technical replicates.

Earlier studies proposed that temperature sensitivity might correlate with subcellular localization shifts, such as nuclear relocalization at elevated temperature (*26*). In contrast, we observed no temperature-dependent changes in the distribution of either Ry_sto_ variant (Suppl. Fig. S4).

At permissive temperature, both WT Ry_sto_ and Ry_sto-K624E_ formed oligomeric complexes >720 kDa upon PVY-CP activation (Fig. 3G). At 30°C, only the WT maintained oligomerization, while the K624E mutant migrated as a 130-kDa monomer (Fig. 3G). Both forms retained AVR binding across temperatures (Fig. 3G), confirming that heat impairs oligomerization rather than recognition.

To assess downstream immunity, we expressed both Ry_sto_ variants in *N. benthamiana*. As expected, both triggered HR when challenged with PVY CP at 22°C, but only WT Ry_sto_ remained active at 30°C (Suppl. Fig. S5). In Arabidopsis transgenics challenged with TuMV-GFP, both lines resisted infection at 22°C, yet at 30°C only the WT prevented systemic virus spread (Fig. 3H-I). These results confirm *in planta* that maintaining correct WHD–LRR orientation is essential for oligomerization and immunity at elevated temperatures, validating our structural model as predictive for stability engineering.

To further test the model’s bidirectionality, we converted a semi–temperature-sensitive TNL, Roq1, into a temperature-tolerant form. According to our modeling, Roq1 activity decreases as temperature rises. Structural modeling revealed that the Roq1 WHD–LRR stabilization depends on a hydrophobic cluster formed by WHD residues 451, 452, 455, 456, and 459 and residues 581, 582, 603, 604, 626, and 627 of the LRR (Fig. 4A). Substituting A455 with Q or N was predicted to reinforce this cluster and add potential hydrogen bonding with T626 (Fig. 4A). We co-expressed native Roq1 and its A455Q and A455N variants, all tagged with Flag, with XopQ-Strep (AVR) in *N. benthamiana roq1* knockouts at 22°C and 30°C. At 22°C, Roq1_A455N_ showed slightly enhanced HR, while Roq1_A455Q_ matched the wild type (Fig. 4 B-D). At elevated temperature, WT Roq1 exhibited reduced HR and oligomerization, whereas both engineered variants remained active, with Roq1_A455N_ showing the strongest response (Fig. 4E-F). BNP confirmed that the TT variants showed a similar pattern of oligomerisation at permissive temperature but maintained significantly higher oligomerization at 30°C compared with wild type (Fig. 4 H-I). Thus, both gain-of-function and loss-of-function experiments validate our analysis of the sequence requirements for temperature-tolerant NLRs.

**Fig. 4.**
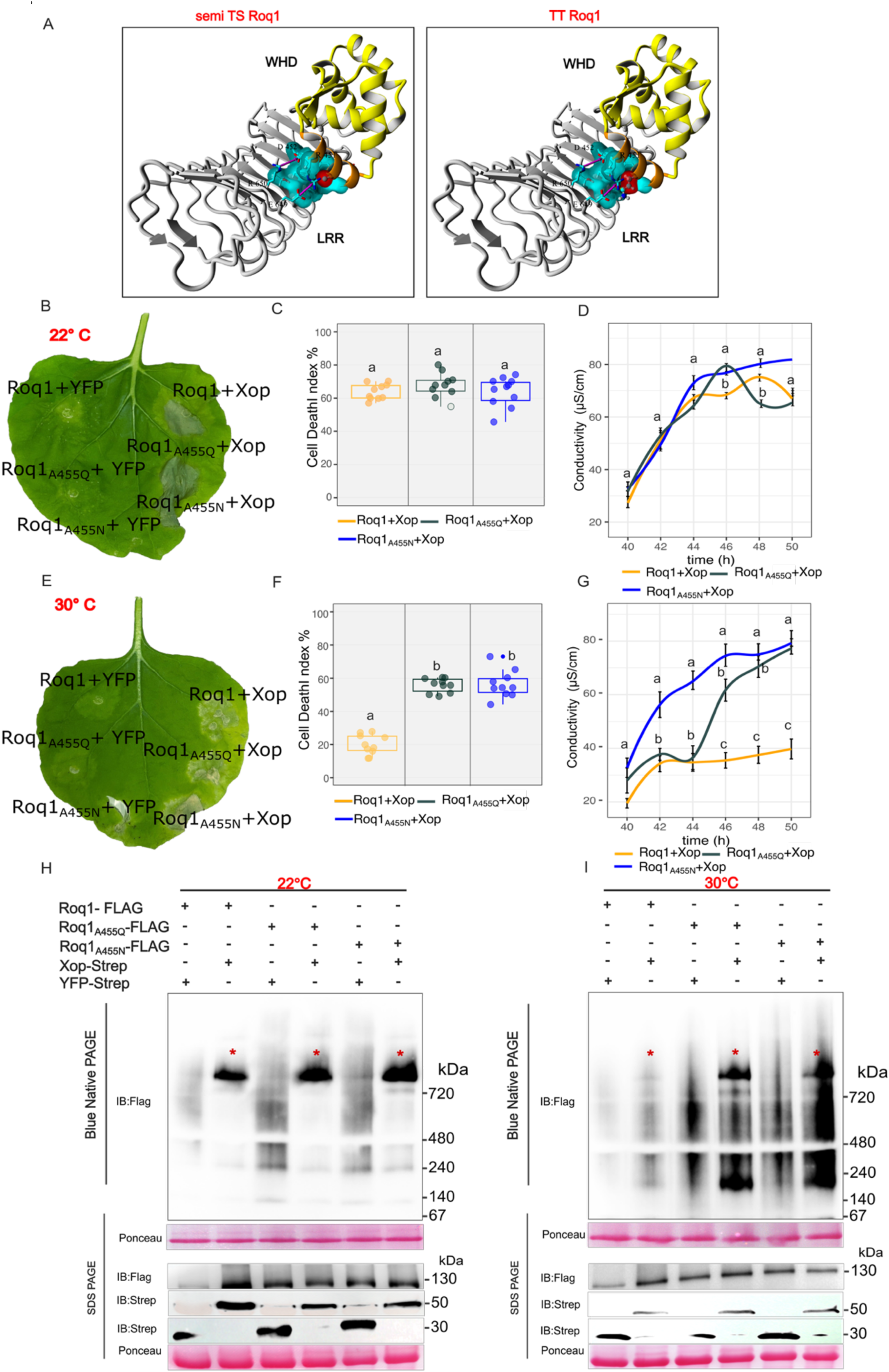
Temperature tolerance of Roq1 increases with enhancing its oligomerization by strengthening WHD-LRR interaction. (A) Left, zoomed-in view of the predicted WHD (yellow) and LRR (gray) regions of Roq1, highlighting the crucial local tertiary interactions; right, Roq1_A455Q_ and Roq1_A455N_ variants predicted to be temperature tolerant. The interacting third helix of the LRR is marked in gold, the cluster of hydrophobic interactions is marked in blue, and the improved region is marked in red. The crucial residues are named and shown as stick or ball-and-stick (when mutated) cartoons. Green balls – carbon, red – oxygen, blue – nitrogen atoms. Magenta arrows – favorable electrostatic interactions. Confidence predictions for the model are provided in Suppl. Table 2. (B-G) Experimental validation of increased temperature tolerance of Roq1 mutant variants. WT Roq1-Flag and its TT variants were co-infiltrated with XopQ-Strep into *roq1*-knockout *N. benthamiana* plants at permissive (22°C; B-D) or elevated (30°C; E-G) temperature. Representative HR phenotypes (B, E), HR scoring box plots (C, F), and ion leakage measurements (D, G) are shown for each temperature. HR was quantified and documented at 5 dpi. Data information: At least three biological replicates and corresponding technical replicates were tested. (H, I) TT Roq1 variants show stronger oligomerization than wild-type Roq1 at elevated temperatures. Proteins were extracted from *N. benthamiana roq1* knockout plants complemented with WT Roq1-Flag, Roq1_A455Q_-Flag, or Roq1_A455N_-Flag co-expressed with XopQ-Strep or YFP-Strep as a control, at both permissive (H) and elevated (I) temperature. Extracted proteins were loaded onto BNP and then immunoblotted. The same samples were used for SDS-PAGE, followed by immunoblotting. Data information: At least three biological replicates and the corresponding technical replicates were tested.

### 1.5 Helper NLRs are predicted to be temperature tolerant

Plant NLRs operate in signaling networks composed of sensor and helper receptors (*5*). Since both temperature-sensitive (TS) and temperature-tolerant (TT) sensors rely on the same limited set of helpers, and modifying sensors alone can alter temperature profiles, we hypothesized that helper NLRs are more likely to be temperature-tolerant.

In *N. benthamiana*, the helper NLR NRG1 mediates both cell death and resistance (*28, 29*). Structural modeling predicted that WHD–LRR stabilization of NRG1 depends on three charge–charge interactions between R638 (LRR) and E470/D466 (WHD) and E642 (LRR), reinforced by stacking between H457/H460 (WHD) and E594 (LRR) (Fig. 5A). To test this, we generated YFP-tagged NRG1 mutants (R638N and R638Q) predicted to disrupt these interactions. NRG1, as part of the signalling TNL pathway, must be co-expressed with a sensor TNL exposed to AVR to test its function. Therefore, to test predicted TS NRG1 variants, we transiently co-expressed them with Ry-Flag and CP-Strep in *nrg1* knockout plants at 22°C and 30°C. At 22°C, all variants triggered comparable hypersensitive responses (HR) and ion leakage (Fig. 5B–D). At 30°C, HR was abolished in R638N and reduced in R638Q, indicating that R638 is critical for maintaining function under heat stress (Fig. 5F–H). Consistent with this phenotype, BNP showed that only WT NRG1 formed oligomers at elevated temperature (Fig. 5J). We also modelled Arabidopsis NRG1.1 and NRG1.2, both showing similar stabilization of WHD–LRR contacts, suggesting conserved temperature tolerance. Because Rx-mediated immunity remains effective at elevated temperature (*30*), we further evaluated WHD–LRR models of *Solanaceae* helpers NRC2, NRC3, and NRC4 (*31*). Each displayed multiple stabilizing electrostatic interactions consistent with TT status. Transient co-expression of Rx and its effector PVX-CP in *nrc2,3,4* knockout plants confirmed that complementation with any NRC restored HR even at 30°C (Suppl. Fig. 6). Together, these results indicate that helper NLRs such as NRG1 and NRCs possess intrinsically stable WHD–LRR interfaces that enable oligomerization and signaling at elevated temperatures, ensuring the robustness of the NLR immune network.

**Fig. 5.**
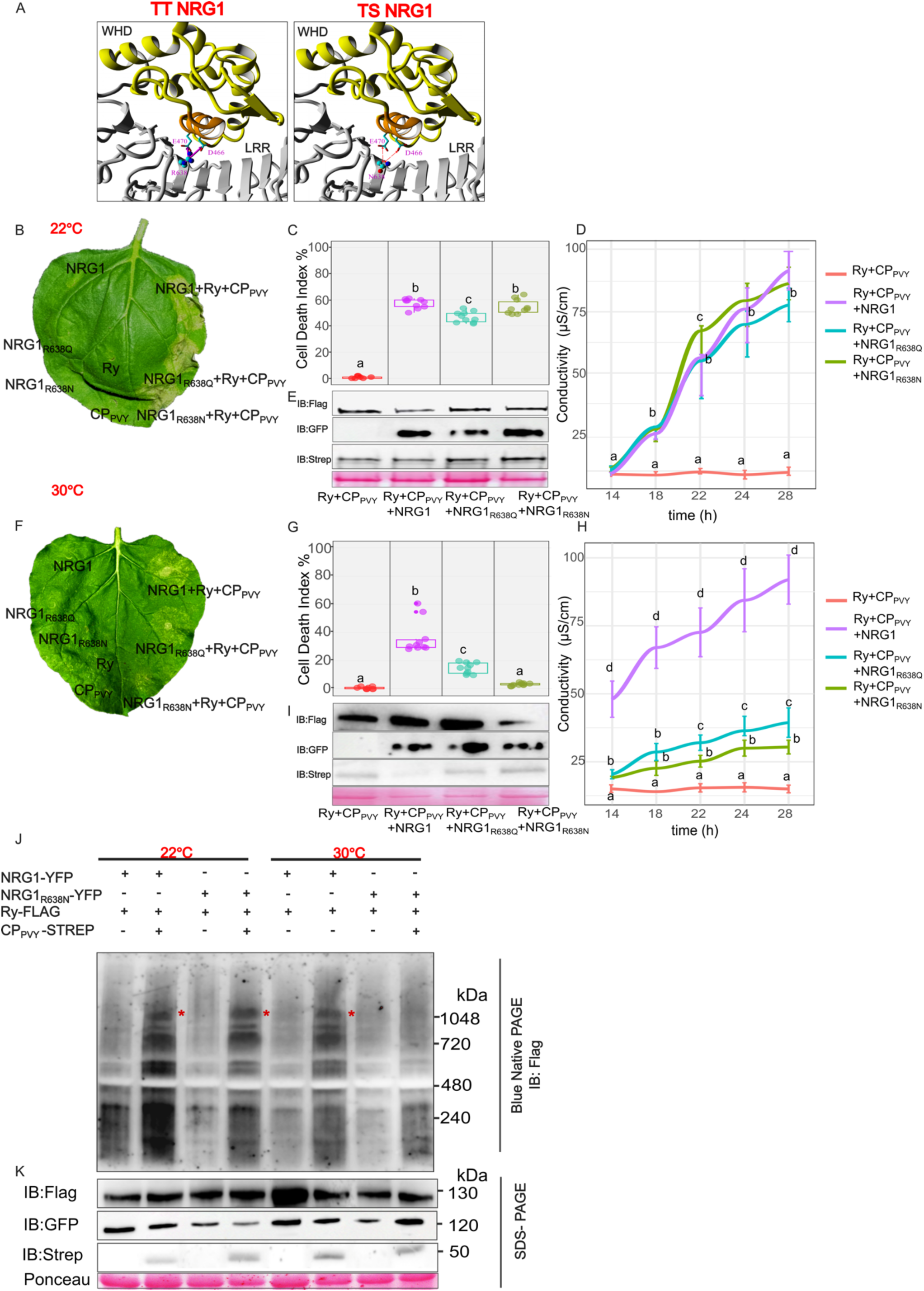
Helper NLR NRG1 is predicted and validated as temperature tolerant. (A) Left, zoomed-in view of the predicted WHD (yellow) and LRR (gray) region of NRG1, highlighting the crucial local tertiary interactions; right, the R638N mutation predicted to turn TT NRG1 into a TS variant (partial phenotype variant R638Q not shown). The interacting third helix of LRR is marked with gold. The crucial residues are named and shown as stick or ball-and-stick (when mutated) cartoons. Green balls – carbon, red – oxygen, blue – nitrogen atoms. Magenta arrows – favorable electrostatic interactions (salt bridges and hydrogen bonds), red arrows show the unfavorable ones, and green arrows represent hydrophobic contacts. Confidence predictions for the model are provided in Suppl. Table 2. (B-I) Experimental validation of temperature tolerance of NRG1. NbNRG1 and its mutant variants were co-expressed with Ry_sto_ and PVY-CP in *nrg1* knockout *N. benthamiana* leaves at permissive (B-E) or elevated (F-I) temperature. Representative HR phenotypes (B, F), HR scoring box plots (C, G), ion leakage measurements (D, H), and protein expression levels (E, I) are shown for each temperature. (J) TS NRG1_R638N_ variant does not form oligomers at elevated temperatures. Proteins were extracted from *N. benthamiana nrg1* knockout plants complemented with WT NRG1 or NRG1_R638N_ variant, co-expressed with Ry_sto_ and PVY-CP or Ry_sto_ only as a control, at both permissive and elevated temperatures. Extracted proteins were loaded on BNP, followed by immunoblotting. The same samples were used for SDS-PAGE, followed by immunoblotting. Ponceau S staining served as a loading control. Data information: For all experiments, a minimum of ten biological repeats, along with the corresponding technical replicates, were performed.

## Discussion

Temperature sensitivity of NLRs has been extensively studied, yet no universal mechanism has been proposed. Here we show that temperature tolerance broadly depends on the efficiency of NLR oligomerization, governed by the orientation and stabilization of the WHD region. Using structural modeling, we classified NLRs by their temperature tolerance and demonstrated that these properties can be rationally modified.

In *Ry_sto_*, disruption of WHD–LRR contacts by the K624E substitution abolishes oligomerization and immune activation at elevated temperature, confirming that structural instability underpins temperature sensitivity. Conversely, engineered Roq1 variants (A455Q, A455N) with strengthened WHD–LRR interactions showed enhanced activity and oligomerization at elevated temperature, demonstrating that temperature resilience can be restored through targeted stabilization. Also, the mutation previously reported for N and SNC1 aligns with our model (*26*), further validating its predictive accuracy.

The subcellular localization of Ry_sto_ and its TS variant remained unchanged across temperatures, contrasting with earlier reports (*26, 32*) of heat-induced nuclear re-localization of NLRs. Comparable stability was observed for Sr6, Sr13, and Sr21 (*33*). Modeling of Sr6 revealed that, although the WHD is stable, disruption of LRR hydrophobic packing, normally reinforced by the conserved axxaxxa/LxxL/IxaxxCxaxxaxx motif, renders the receptor heat-sensitive. Many NLRs depend on helper NLRs for immune signaling (*5, 34*), and Rx1-interacting NRCs remain active at elevated temperatures (*30*). Consistent with this, *in silico* analysis predicted that WHD regions of NRC1–4 and NRG1 are intrinsically stable, and a single mutation converting NRG1 to a TS form experimentally confirmed this tolerance. Together, these findings establish a mechanistic and predictive framework for engineering durable NLR function under fluctuating environmental conditions.

## Acknowledgements

We thank Prof. Jane Parker for sharing seeds of *N. benthamiana eds1* knockout plants, Prof. James Carrington for sharing the TuMV-GFP infectious clone, and Prof. Sophien Kamoun for *nrc2* and *3* constructs and *nrc2,3,4* knockout plants seeds. We also thank Prof Dangl for critical reading of the manuscript.

The research was supported by the OPUS grant nr 2022/47/B/NZ9/02224 from the National Science Centre, Poland, awarded to M. G-B. We are also grateful to 2Blades for support.

## Author contributions

Conceptualization: M.G.-B., J.T.P., K.W., J.H. Methodology: M.G.-B., J.T.P., H.-K.A. Formal analysis: M.G.-B., K.W. Investigation: M.G.-B., J.T.P., H.-K.A., T.V.-C., M.L., I.B.-F., K.W., J.H. Writing – Original Draft: M.G.-B., K.W., A.I.W., J.T.P., J.D.G.J. Writing – Review & Editing: all authors. Visualization: M.G.-B., J.T.P., M.L. Supervision: M.G.-B.

## Data Availability

Reagents, tools, and materials generated in this study are available from the corresponding author upon request.

## Materials and Methods

### Plant material

*N. benthamiana* wild-type (WT) *roq1, eds-1*, *nrg1, and nrc2,3,4 Nicotiana* knockouts were grown for 6 weeks in soil under controlled environmental conditions (16 h light / 8 h dark) under permissive (22°C) and elevated (28°C to 30°C) temperatures as described previously (*35*). Transgenic *N. benthamiana* plants were obtained as described (*23*). Transgenic *Arabidopsis* plants (ecotype Columbia) were obtained using the floral dip method as described (*36*). Plants were then cultivated in Jiffy7 pots in controlled-environment chambers (Percival Scientific, Perry, IA, USA) at 22°C or 30°C with 40% humidity, under 8 h of light. All details are shown in Suppl. Table 3.

### Constructs assembly

*Ry_sto_* and *Roq1* native form, together with their TS and TT variants (obtained through site-directed mutagenesis), *PVY-CP*, and *N* constructs were assembled using the Golden Gate Modular Cloning (MoClo) system (*37*), along with the MoClo Plant Parts Kit (*38*). To prevent NRG1 autoactivity, the estradiol-inducible destination vector was used for NRG1 expression. *NbNRG1* cDNA was PCR-amplified and cloned into the entry pENTR/D-TOPO vector. The resulting entry clones were LR recombined with the Gateway pMDC7 containing a C-terminal YFP-HA, as described (*39*). A similar approach was used for the TS NRG1 variants tested, which were obtained via site-directed mutagenesis. The constructs are listed in the Suppl. Table 3.

### Structural modeling

Structures of the receptors were initially modeled by homology with the local instance of the Yasara Structure package (v. 21.8.27). Up to 8 template structures were analyzed; for each, up to 5 alignment variations were tested. For each modeled loop, up to 50 conformations were tested. The maximal oligomerization state was limited to the tetramer. The quality of the resulting models was assessed with the Z-score value, which distribution along the sequence was monitored to identify the questionable regions of the protein. In most cases, a model built on the EM structure of a disease resistance protein (i.e., 7CRC, 7DFV, 7CRB, 7JLV, or 7JLU) was scored as the best, with a preference for the tetrameric structures build on a shorter template (7DFV), which still covered the regions of interest (WHD and LRR domains). Finally, the best parts of the single-template models were combined into a hybrid model.

The active form of any tested receptor was always assigned to the best-scored tetramer.

Screening of the receptor structures was performed using the local instance of the AlphaFold v2.3.2 package. We have used either Monomer or Multimer versions to model a monomeric or tetrameric structure of a target protein. After identifying the interaction pattern, which is crucial for stability prediction, the monomeric form of the receptor was modeled. The quality of the predicted structures was analyzed using the residue-specific pLDDT (predicted Local Distance Difference Test), and the mean of the backbone pLDDTs was treated as the descriptor of the overall structure quality. In parallel, we have also calculated the predicted template modelling (pTM) score, which value varied in the range of 0.58-0.87. For the region of interest covering 365 residues located in the WHD domain and the N-terminal fragment of the LRR domain, the pTM values were significantly higher (0.81–0.92), indicating high-quality predictions. All data are shown in Suppl. Table 2.

### Transient expression assay

Binary constructs for transient expression were transformed into *Agrobacterium tumefaciens* strain GV3101 (pMP90). *Agrobacterium* cells were resuspended in an infiltration buffer (10 mM MgCl₂ and 10 mM MES, pH 5.6, 100 μM acetosyringone). The optical density at 600 nm (OD₆₀₀) of each construct-containing *Agrobacterium* suspension was adjusted to 0.5, except for *Agrobacterium* containing p19, where it was adjusted to 0.2. Infiltration suspensions were incubated for 1-2 hours at room temperature prior to infiltration. Hypersensitive response (HR)-related phenotypes were evaluated at three days post-infiltration (dpi).

### HR scoring

For the HR assay, 4-week-old *N. benthamiana* plants were used. Constructs in *Agrobacterium* were infiltrated or co-infiltrated into the abaxial surface of *N. benthamiana* leaves. The HR phenotypes were scored at 5 dpi. Detached leaves were photographed under UV light with a yellow filter (Wratten K2 Yellow Filter no. 8; Kodak) attached to the camera lens. Images were captured with a 2-second-long exposure, F8.0, and ISO 400, then scored using ImageJ software according to the scale described in Grech-Baran et al., 2022 (*24*).

### Ion conductivity

At the specified time points, eight leaf discs (1 cm in diameter) were cut from infiltrated zones and floated abaxial side up on 5 ml of MilliQ water for 10 minutes at 18°C with gentle stirring (50 rpm). The conductivity of the water was measured using a WTW InoLab Multi 9310 IDSCDM83 benchtop meter and reported in μS cm^−1^.

### Confocal laser scanning microscopy

Subcellular localization of the fusion proteins was examined using a Nikon C1 confocal system built on TE2000E and equipped with a 60x Plan-Apochromat oil immersion objective (Nikon Instruments BV Europe, Amsterdam, the Netherlands). GFP was excited by the Sapphire 488 nm laser (Coherent, Santa Clara, CA, USA) and observed with the 515/530 nm emission filter, while RFP was excited by the 543 nm helium–neon laser and detected with the 605/75 nm barrier filter. GFP and RFP imaging were performed sequentially to prevent bleed-through. Fluorescence microscopy was conducted at the Fluorescence Microscopy Facility of IBB PAS, Poland. Confocal images were analyzed using the free viewer EZ-C1 and ImageJ software.

### Protein extraction

For protein extraction in Figs 1, 3, and 4, samples were collected from *Agrobacterium*-infiltrated leaves at 2 dpi using a cork borer. All leaf discs were transferred into 2 ml Eppendorf tubes and rapidly frozen in liquid nitrogen. The samples were then ground into a powder. A protein extraction buffer containing 50 mM Tris-Cl (pH 7.5), 150 mM NaCl, 10% glycerol, 1 mM EDTA pH 8.0, 5 mM DTT, 0.2% IGEPAL, and protease inhibitor cocktail was vortexed with the ground tissue. Centrifugation at 18,000 g for 15 minutes, followed by 5 minutes, was used to remove cell debris. Aliquots of the samples were either directly used for Blue Native-PAGE (BNP) or subjected to immunoprecipitation (IP) on Flag beads, then eluted with 3xFlag peptide and loaded onto BNP. For SDS-PAGE, aliquots of the input and Co-IP samples were mixed with 3X SDS sample buffer (containing 30% glycerol, 3% SDS, 93.75 mM Tris-Cl pH 6.8, and 0.06% bromophenol blue) with 100 mM DTT and heated at 70°C for 10 minutes.

To assess protein expression levels following the physical assays (HR-scoring and ion-leakage), lysates were prepared with the same buffer as used for BNP and then mixed with 3× SDS sample buffer for SDS-PAGE. Subsequently, immunoblotting with an appropriate antibody was performed.

For Fig. 5, leaves were infiltrated with *Agrobacterium* containing each construct at a ratio of 1:1:3 (RY: CP: NbNRG1). Notably, NRG1 was induced 32 hours after infiltration by spraying with 100 µM β-estradiol (Sigma-Aldrich; St. Louis, USA) and 0.002% (v/v) Silwet L-77. Tissue samples for protein assays were collected within 16 hours post-induction. All leaf discs were placed in 2-ml Eppendorf tubes, frozen in liquid nitrogen, and ground. Protein extraction buffer (50 mM Tris-Cl, pH 7.5, 150 mM NaCl, 10% glycerol, 1 mM EDTA, 5 mM DTT, 0.1% DDM, protease inhibitor cocktail) was vortexed with the ground tissue. Centrifugation was performed at 23,000 g for 30 minutes to remove cell debris. Samples were aliquoted and used directly for BNP and for SDS-PAGE.

To assess protein levels after the physical assays (as mentioned above), lysates were prepared using an extraction buffer (100 mM Tris-Cl pH 7.5, 150 mM NaCl, 0,2% IGEPAL, 1 mM EDTA pH 8.0, 0.1% SDS, 10 mM DTT, and protease inhibitor cocktail). The lysates were then mixed with 3X SDS sample buffer for SDS-PAGE, followed by immunoblotting with an appropriate antibody.

### Immunoprecipitation and elution

Flag-M2 beads (Sigma, A2220) were added to the protein extract after centrifugation and incubated at 4°C for 1 hour. Afterward, the beads were washed five times with protein extraction buffer. For protein elution, 3xFlag peptide (Sigma, F4799) was added to the beads at a concentration of 0.2 mg/ml and incubated for 1 hour. Sample aliquots were prepared for Blue Native PAGE (BNP) analysis and immediately frozen in liquid nitrogen.

### BNP and SDS-PAGE analysis

For blue native polyacrylamide gel electrophoresis (BNP), clarified protein extracts were diluted according to the manufacturer’s instructions with NativePAGE™ 5% G-250 sample additive, 4× NativePAGE Sample Buffer, and deionized water. Samples were loaded onto 3%–12% Bis-Tris NativePAGE™ gels (Invitrogen) and ran alongside SERVA Native Marker (SERVA). Following the separation, proteins were transferred to polyvinylidene difluoride (PVDF) membranes using NuPAGE™ Transfer Buffer and the Trans-Blot Turbo Transfer System (Bio-Rad), according to the manufacturer’s protocol. Membranes were fixed with 8% acetic acid for 15 minutes, rinsed with water, and air-dried. To visualize native protein standards, membranes were reactivated with ethanol prior to immunoblotting. For SDS-PAGE, protein samples were mixed with SDS loading buffer and denatured at 72°C for 10 minutes. Proteins were transferred to PVDF membranes using the Trans-Blot Turbo Transfer System with the supplied transfer buffer, following standard procedures.

### Immunoblotting and detection

Membranes were blocked in 5% (w/v) non-fat dry milk in Tris-buffered saline containing 0.01% Tween-20 (TBS-T) for 1 hour at room temperature. Blots were then incubated overnight at 4°C with horseradish peroxidase (HRP)-conjugated primary antibodies diluted 1:5,000 in 5% milk / TBS-T. The following antibodies were used: anti-GFP (B-2, Santa Cruz Biotechnology), anti-Strep (71591-M, Sigma-Aldrich), anti-Flag (M2, Sigma-Aldrich), and anti-Myc (conjugated HRP, Merck clone 9E10). Signal detection was performed using Pierce™ ECL Western Blotting Substrate (32106, Thermo Fisher Scientific), with up to 50% SuperSignal™ West Femto Maximum Sensitivity Substrate (34095, Thermo Fisher Scientific) added for enhanced sensitivity when needed. Chemiluminescence was imaged using Azure 400 AZI400-01 (Azure Biosystems, USA). Membrane loading controls were visualized by staining with Ponceau S (Sigma-Aldrich).

### Viral Infectious Assay

For the TuMV-GFP infection, plants were inoculated using the TuMV-GFP infectious clone as described by Garcia-Ruiz et al., 2015 (*40*).

### Statistical analysis

Statistical analyses were performed using R 4.5.0 within R Studio 2025.05.0+496. Technical replicates included multiple readings from the same plant in a single experiment, while biological replicates involved measurements from independent plants. Data analysis pipeline was as follows: first, data were tested for suitability for parametric analysis by assessing the normality of residuals with the Shapiro–Wilk test. If the data met the assumptions for parametric testing, a repeated-measures analysis of variance (ANOVA) was conducted, followed by Tukey’s honestly significant difference (HSD) test (**, P < 0.001).

---

**Suppl. Fig. 1.**
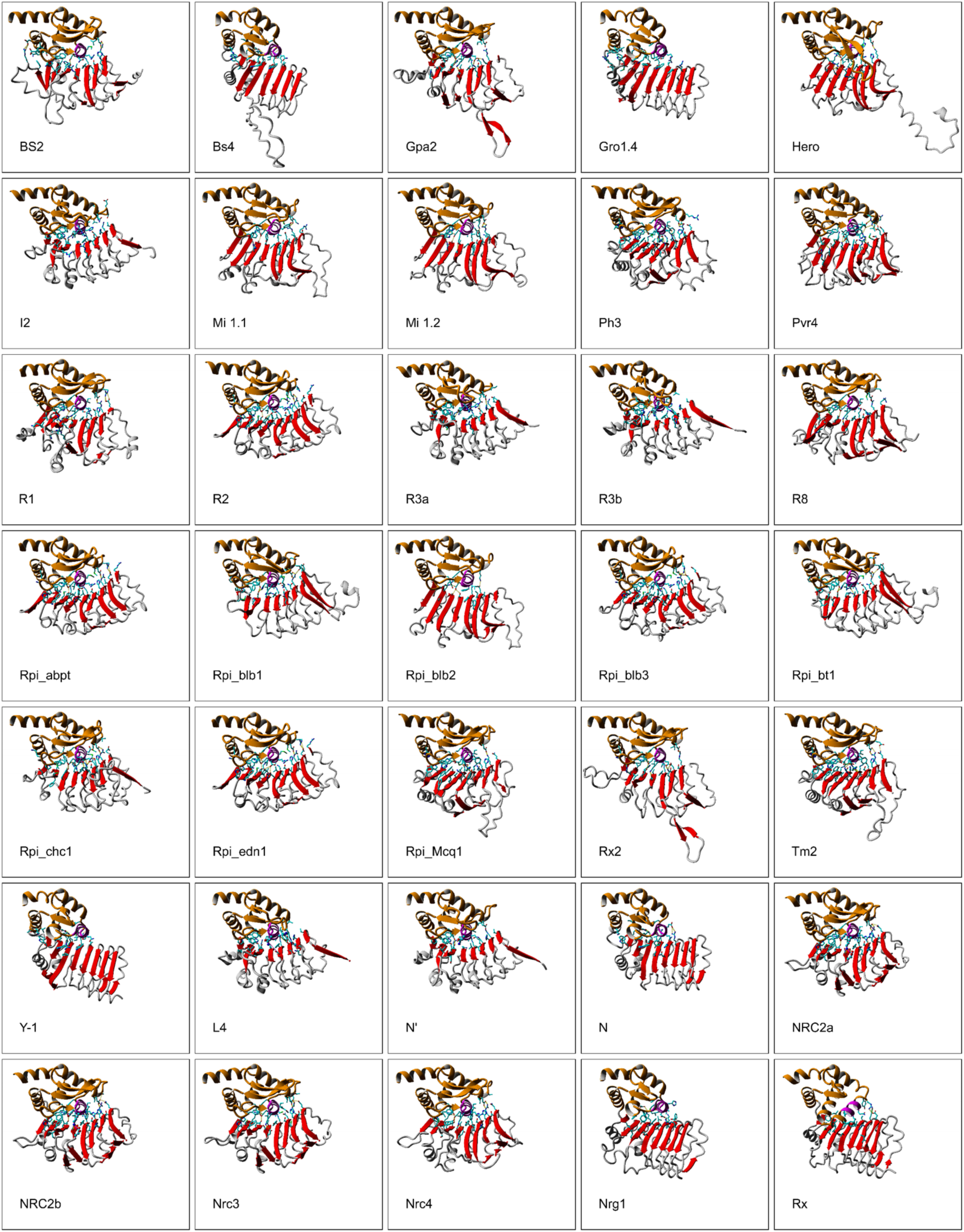
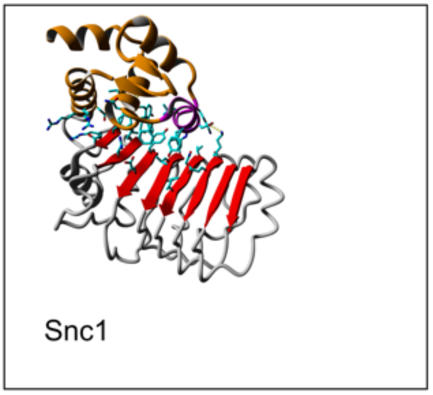
Structural models of the WHDs and interacting LRR regions of Ry_sto_ and various other *Solanaceae* receptors with known and unknown temperature sensitivity. The third helix, crucial for WHD-LRR interaction, is highlighted in magenta within the WHD region (gold). Side chains of residues involved in the surface contacts with LRR are also shown. The beta-sheet of the LRR domain is marked in red. The receptor structures were screened using the AlphaFold v2.3.2 package. The predicted pTM score ranged from 0.81 to 0.92 (Suppl. Table 2). Models were visualized using Yasara (Krieger et al., 2009).

**Suppl. Fig. 2.**
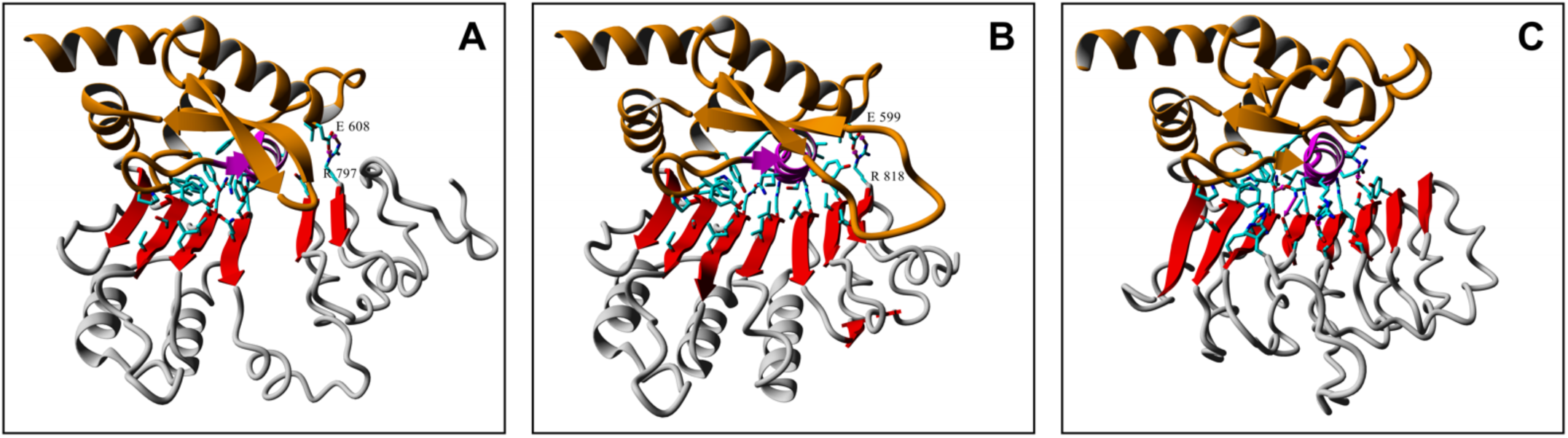
Structural models of the WHD-LRR regions for Rx (A), Sr21 (B), and Mi-1.1 (C). The third helix, crucial for WHD-LRR interaction, is highlighted in magenta within the WHD region (gold). Side chains of residues involved in the surface contacts with LRR are also shown. The beta-sheet of the LRR domain is marked in red. The receptor structures were screened using AlphaFold v2.3.2 package. The predicted pTM score for all models was > 0.9 (Suppl. Table 2). Models were visualized using Yasara (Krieger et al., 2009).

**Suppl. Fig. 3.**
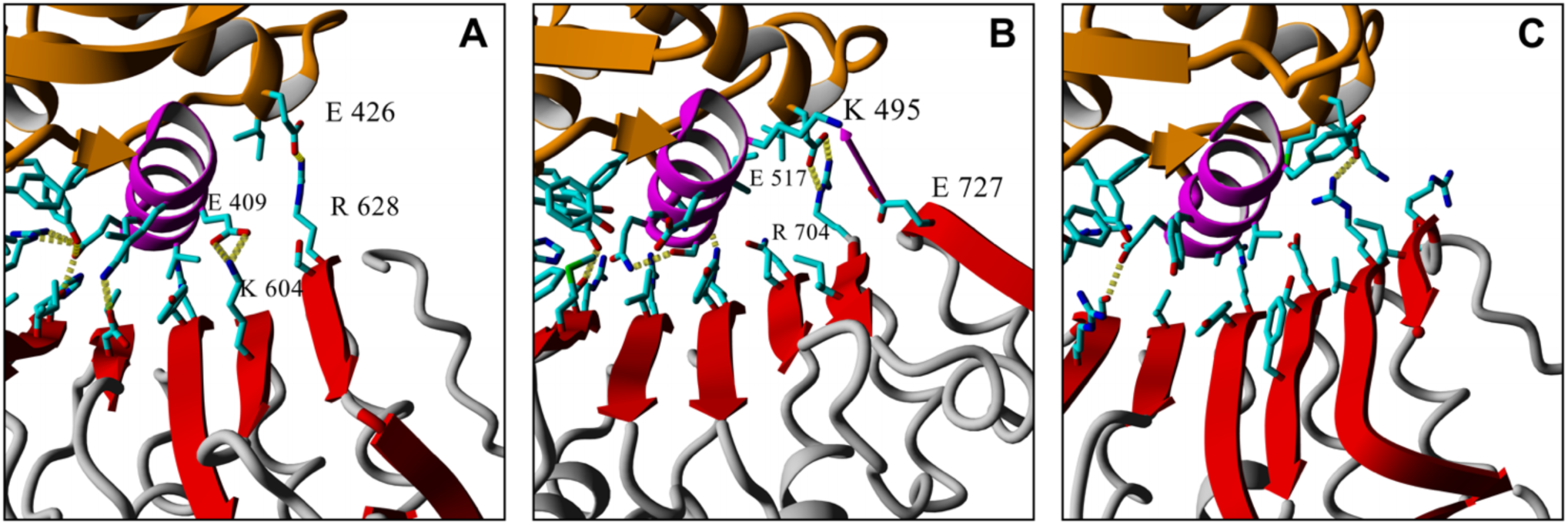
Structural models of the WHD-LRR regions for Yr5a (A), Yr7 (B), and Yr87/Lr85 (C) immune receptors. The third helix, crucial for WHD-LRR interaction, is highlighted in magenta within the WHD region (gold). Side chains of residues involved in the surface contacts with LRR are also shown. The beta-sheet of the LRR domain is marked in red. The receptor structures were screened using AlphaFold v2.3.2 package. The predicted pTM score for all models was > 0.8 (Suppl. Table 2). Models were visualized using Yasara (Krieger et al., 2009).

**Suppl. Fig. 4.**
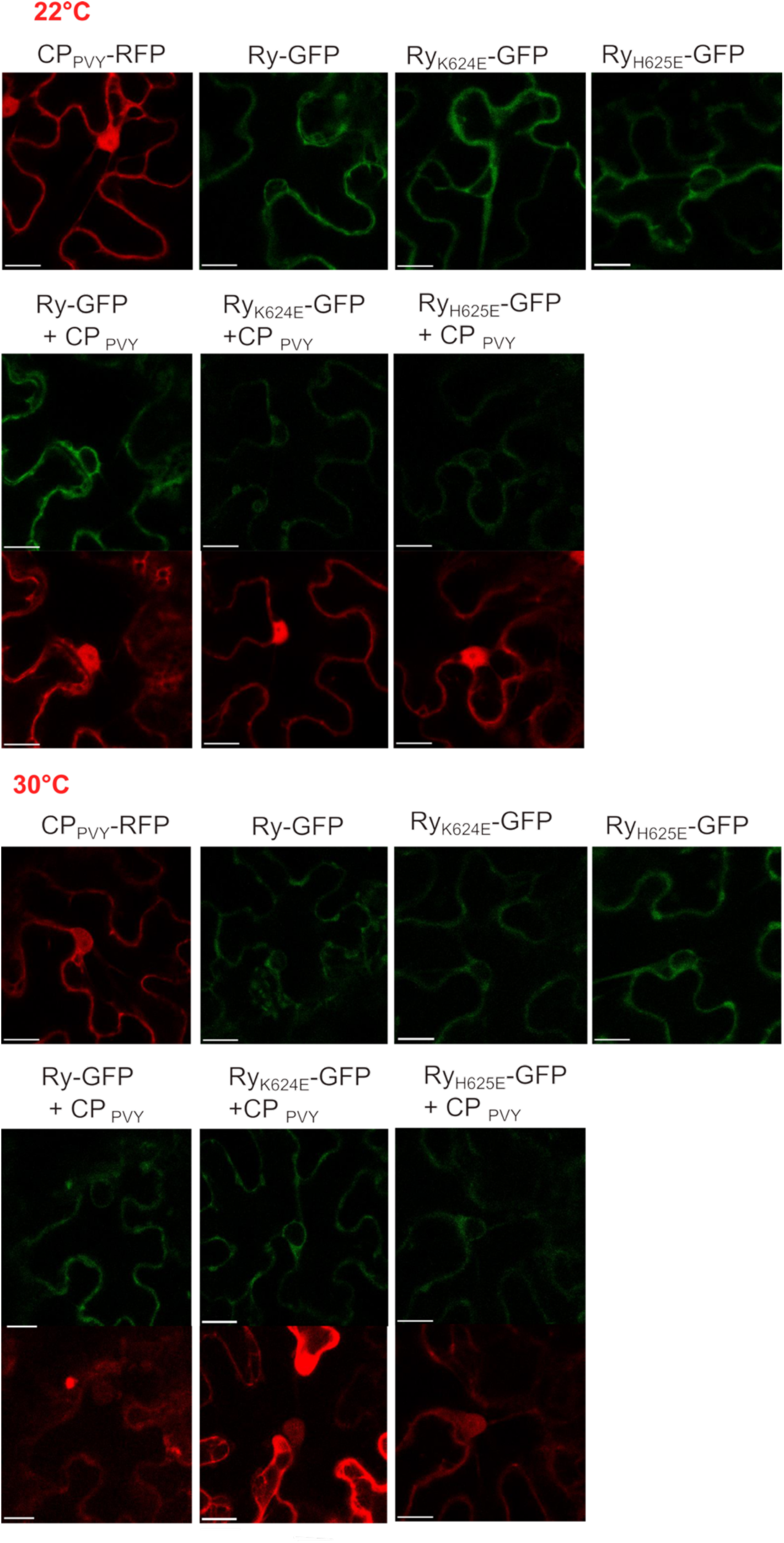
The subcellular localization of Ry_sto_ and its temperature-sensitive variant does not change with temperature increase. Cellular localization of Ry_sto_ and its variants at permissive (22°C) and elevated (30°C) temperatures in representative *N. benthamiana* leaf epidermal cells. Cells were transiently co-expressing Ry_sto_ or its mutant variants with PVY-CP. Confocal images were captured at 2 dpi. Approximately 50 transformed cells were analyzed per variant. Bar, 10 µm.

**Suppl. Fig. 5.**
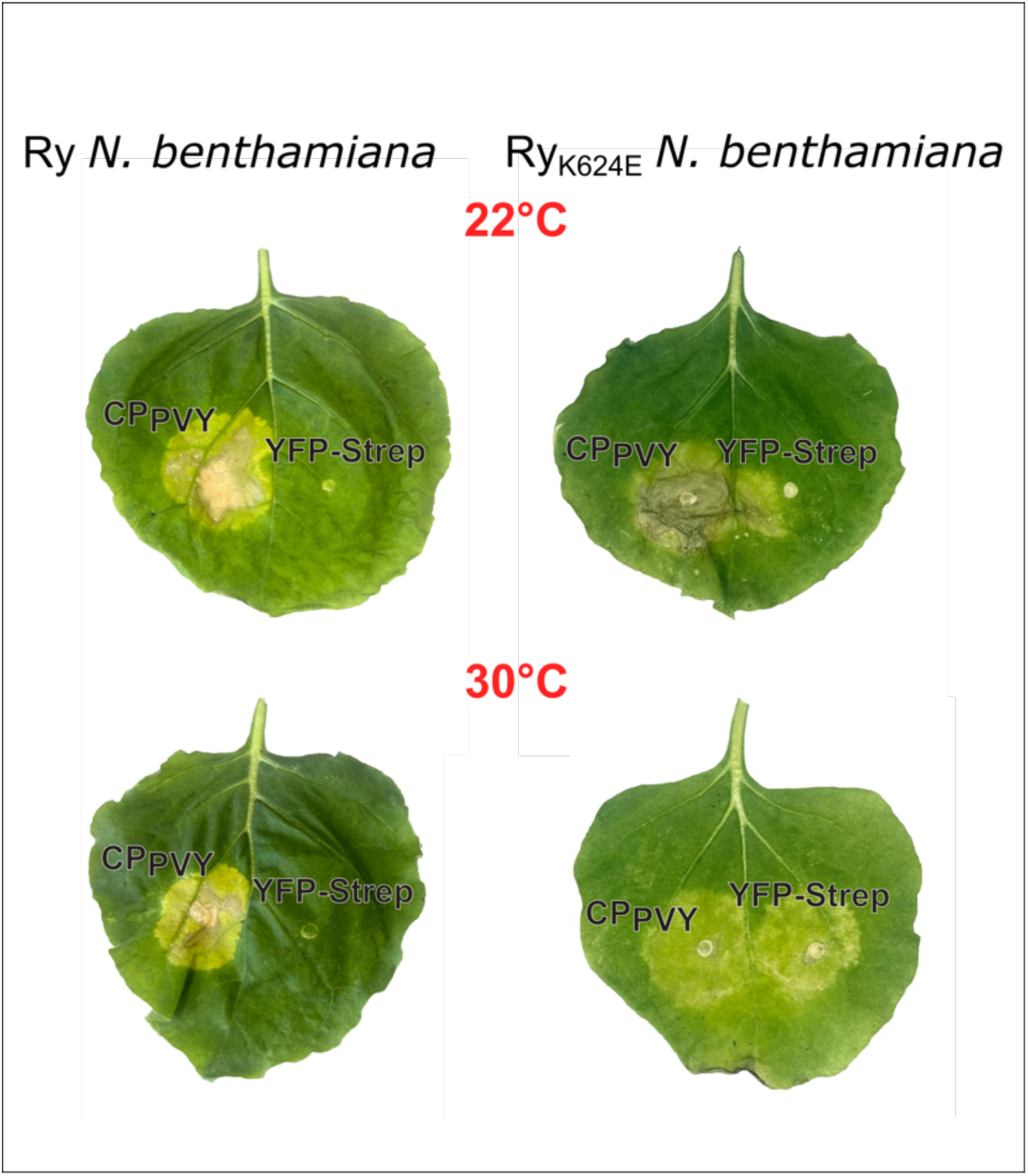
Abolition of HR in TS Ry_sto-K624E_ transgenic lines at elevated temperature. Transgenic Ry_sto_ and Ry_sto-K624E_ *N. benthamiana* plant lines were agroinfiltrated with PVY-CP or YFP as a control at both permissive (22°C) and elevated (30°C) temperature. An HR reaction was observed in both plant genotypes at the permissive temperature (A). At the elevated temperature (B), only lines expressing the native Ry_sto_ variant could activate the HR. The experiment was conducted on five different lines for each genotype. A representative photograph of the leaves was taken at 5 dpi.

**Suppl. Fig. 6.**
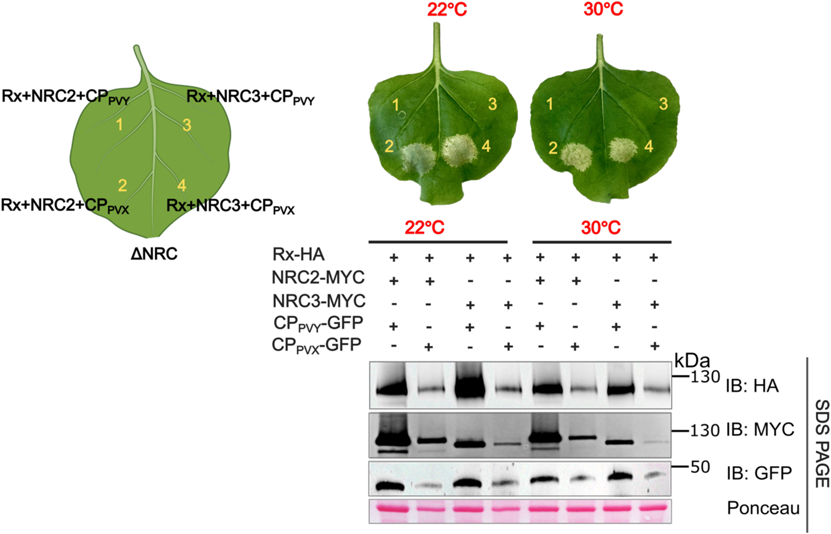
NRCs, CC-NLR helpers in *Solanaceae* are temperature-tolerant. To experimentally verify the predicted temperature-tolerant profile of NRC helper proteins (Suppl. Table 2), we co-expressed a sensor CC-NLR Rx and its AVR PVX-CP along with NRC2 or NRC3, in *nrc 2,3,4* knockout plants at both permissive (22°C) and elevated (30°C) temperature. HR phenotype was observed in both the Rx-PVXCP-NRC2 and Rx-PVXCP-NRC3 co-expression systems at both temperatures.

**Suppl. Fig. 7.**
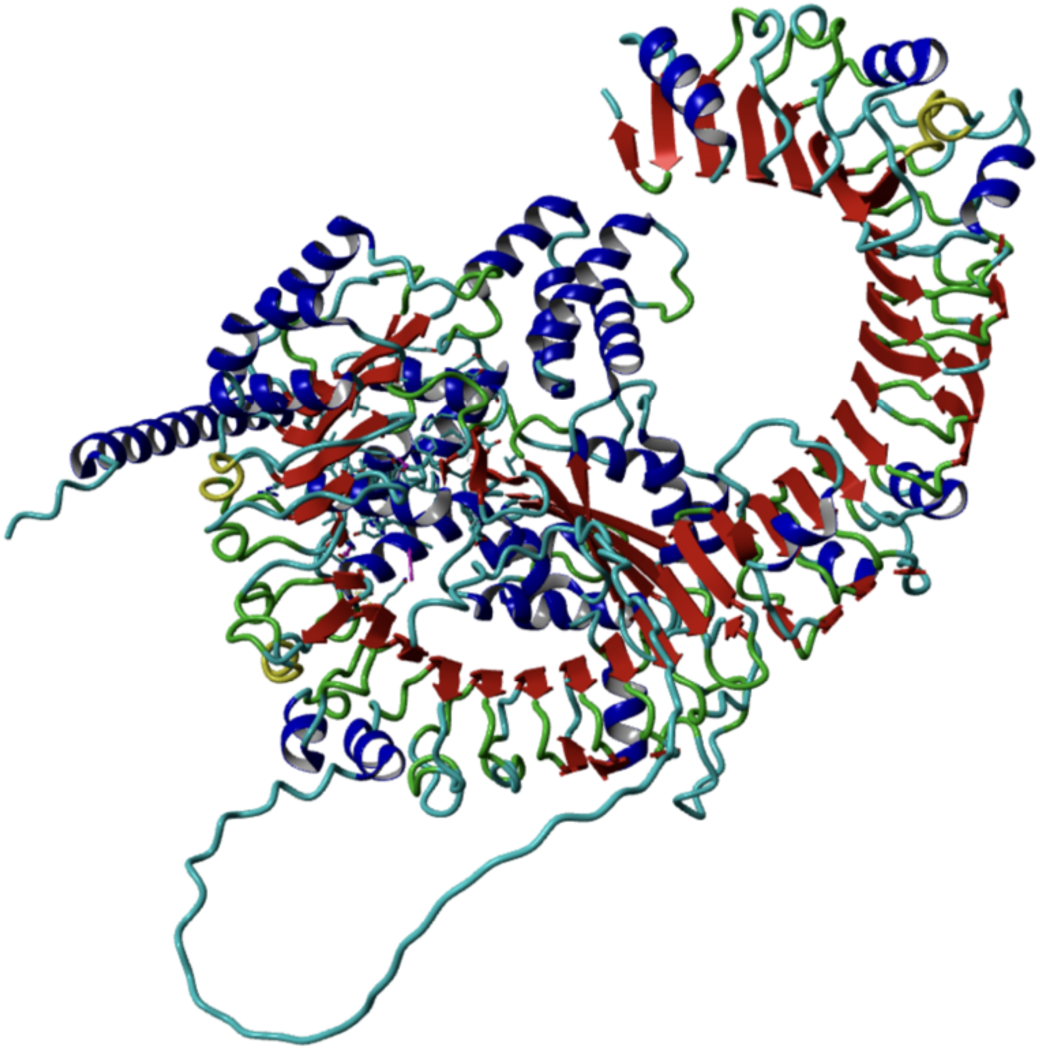
Structural model of Sr6. The model shows that the packing of the LRR domain, which relies on the conserved structural motif axxaxxa/LxxL/IxaxxCxaxxaxx, is disrupted. Instead of multiple hydrophobic interactions supporting the LRR structure, hydrogen bonds mainly stabilize the Sr6 LRR domain. The receptor structure was screened using AlphaFold3. The predicted pTM score =0.7. The model was visualized using Yasara (Krieger et al., 2009).

**Supplementary Table 1.** A list of criteria used to assess temperature sensitivity profiles of the studied receptors. The calculations were made with a 3.3 Å threshold for the distance between heavy atoms.

| RECEPTOR | Number of close pairs of residues located in particular WHD regions and LRR, respectively |  |  |  |  |  |  |  |
| --- | --- | --- | --- | --- | --- | --- | --- | --- |
|  |  |  | H3 (WHD) |  |  |  | H2(WHD) | H4(WHD) |
|  | total | hydrophobic residues | anion-cation | His-anion | anion-anion cation-cation | His-cation | anion-cation | anion-cation |
| BS2 | 12 | 1 | 1 | 1 | 1 | 1 | 0 | 1 |
| Bs4 | 3 | 0 | 0 | 0 | 0 | 0 | 2 | 0 |
| Gpa2 | 9 | 2 | 4 | 1 | 0 | 0 | 0 | 0 |
| Gro1_4 | 6 | 1 | 0 | 0 | 0 | 1 | 2 | 0 |
| Hero | 11 | 2 | 2 | 0 | 0 | 0 | 1 | 1 |
| I2 | 9 | 1 | 0 | 0 | 1 | 1 | 0 | 0 |
| Mi_1_1 | 9 | 2 | 2 | 0 | 0 | 0 | 0 | 0 |
| Mi_1_2 | 14 | 5 | 2 | 0 | 0 | 0 | 0 | 2 |
| Ph3 | 18 | 3 | 1 | 0 | 1 | 0 | 0 | 1 |
| Pvr4 | 13 | 1 | 2 | 0 | 0 | 0 | 1 | 2 |
| R1 | 12 | 1 | 2 | 1 | 0 | 1 | 1 | 2 |
| R2 | 8 | 2 | 1 | 0 | 0 | 0 | 0 | 1 |
| R3a | 10 | 2 | 0 | 0 | 0 | 1 | 1 | 2 |
| R3b | 8 | 2 | 0 | 0 | 0 | 1 | 0 | 1 |
| R8 | 9 | 1 | 2 | 0 | 0 | 0 | 0 | 1 |
| Rpi_abpt | 8 | 2 | 2 | 0 | 0 | 0 | 0 | 2 |
| Rpi_blb1 | 9 | 1 | 0 | 1 | 0 | 0 | 0 | 2 |
| Rpi_blb2 | 7 | 1 | 1 | 0 | 0 | 0 | 0 | 1 |
| Rpi_blb3 | 12 | 2 | 3 | 0 | 0 | 0 | 0 | 2 |
| Rpi_bt1 | 9 | 1 | 0 | 1 | 0 | 0 | 0 | 2 |
| Rpi_chc1 | 18 | 2 | 3 | 0 | 0 | 0 | 0 | 2 |
| Rpi_edn1 | 11 | 3 | 3 | 0 | 0 | 0 | 0 | 2 |
| Rpi_Mcq1 | 16 | 2 | 3 | 0 | 3 | 0 | 0 | 2 |
| Rpi_sto1 | 9 | 1 | 0 | 1 | 0 | 0 | 0 | 2 |
| Rx2 | 7 | 1 | 3 | 1 | 0 | 0 | 0 | 1 |
| Tm_2 | 12 | 2 | 3 | 0 | 2 | 0 | 0 | 2 |
| Y_1 | 6 | 2 | 0 | 0 | 0 | 0 | 1 | 0 |
| L4 | 10 | 2 | 0 | 0 | 0 | 1 | 0 | 0 |
| N' | 9 | 1 | 0 | 0 | 0 | 0 | 0 | 0 |
| N | 11 | 1 | 1 | 0 | 0 | 0 | 2 | 0 |
| NRC2a | 11 | 2 | 3 | 0 | 0 | 1 | 0 | 1 |
| NRC2b | 11 | 2 | 3 | 0 | 0 | 1 | 0 | 1 |
| Nrc3 | 9 | 2 | 3 | 0 | 0 | 0 | 0 | 2 |
| Nrc4 | 12 | 3 | 3 | 0 | 0 | 1 | 0 | 1 |
| Nrg1 | 6 | 4 | 4 | 0 | 0 | 0 | 1 | 0 |
| Roq1 | 9 | 6 | 2 | 0 | 0 | 0 | 1 | 0 |
| Rx | 6 | 1 | 0 | 0 | 0 | 0 | 0 | 0 |
| Snc1 | 11 | 3 | 2 | 0 | 1 | 0 | 1 | 0 |
| Rpi_amr1 | 12 | 2 | 2 | 0 | 0 | 1 | 0 | 2 |
| Rpi_amr3i | 14 | 2 | 2 | 0 | 1 | 0 | 0 | 2 |
| SR21 | 12 | 0 | 0 | 0 | 0 | 1 | 0 | 1 |
| Ry<sub>sto</sub> | 14 | 2 | 3 | 0 | 0 | 0 | 0 | 0 |

colour code: red- TT, blue- semi TT, black- TS

**Supplementary Table 2.**
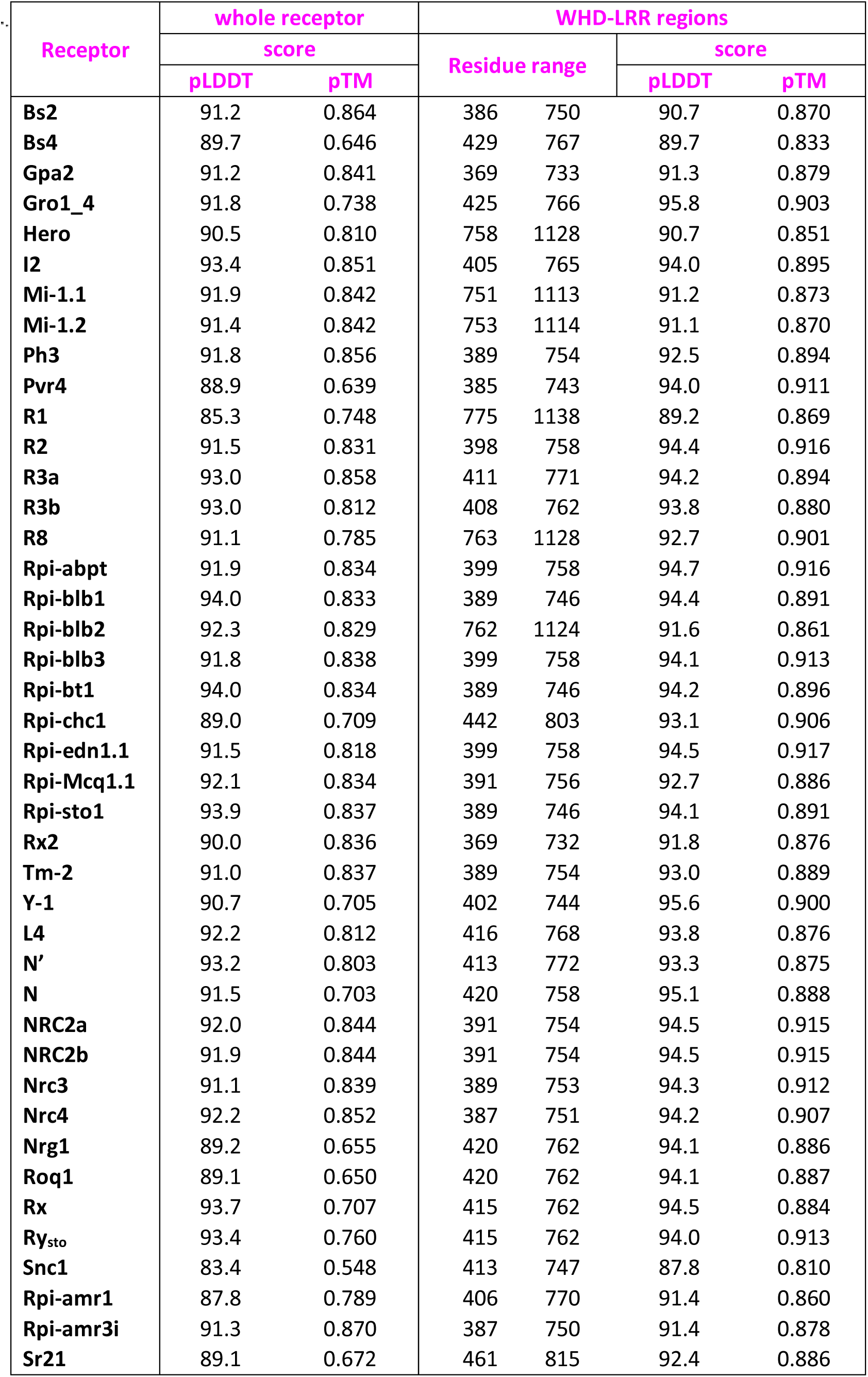
Confidence scores (pLDDT and pTM) assessed for each receptor modeled, supplemented with values calculated separately for the WHD-LRR regions.

**Supplementary Table 3.**
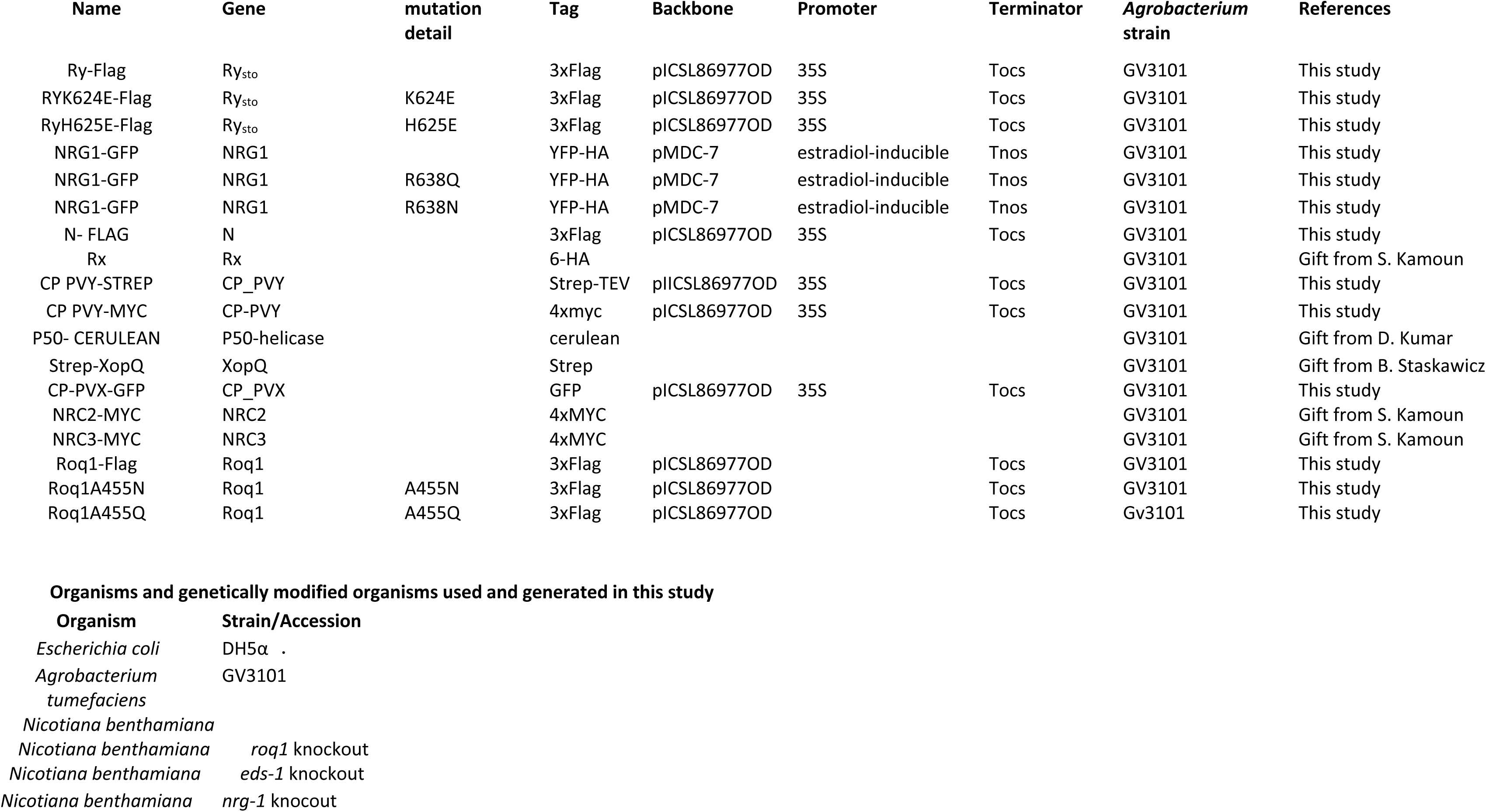
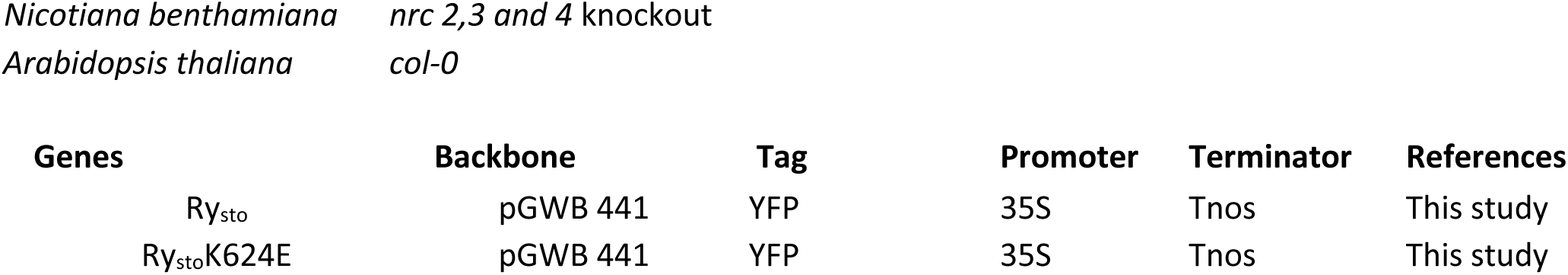
A list of organisms, constructs, and genetically modified organisms used and generated in this study.

